# PNAG exopolysaccharide eradication gives neutrophils access to *Staphylococcus aureus* biofilm infections

**DOI:** 10.1101/2023.01.23.525131

**Authors:** Rachel M. Kratofil, Trevor E. Randall, Josefien W. Hommes, Rehnuma Sejuty, Jessica Chisholm, Deepa Raju, Mario Vargas, P. Lynne Howell, Gerald B. Pier, Douglas W. Morck, Joe J. Harrison, Paul Kubes

## Abstract

*Staphylococcus aureus* (*S. aureus*) can form biofilms on biotic or abiotic surfaces making biofilm infections a relevant clinical problem. Biofilms can evade immunity and resist antimicrobial treatment, and as such an understanding of biofilm infection *in vivo* is necessary to inform new therapeutics. Using a mouse model of *S. aureus* foreign-body skin infection and intravital microscopy, we imaged the interactions between neutrophils and *S. aureus* biofilm. We observed that neutrophils were separated from bacteria by a biofilm matrix composed of the polysaccharide intercellular adhesin (PIA), an exopolysaccharide chemically designated as poly-N-acetylglucosamine (PNAG) that is produced by enzymatic machinery encoded by the *icaADBC* operon. Infection with *icaADBC-deficient S. aureus* strains led to increased neutrophil infiltration and access to bacteria and resulted in full clearance of infection by 7 days. Moreover, enzymatic treatment with PgaB, which hydrolyzes partially deacetylated PNAG, was shown to disaggregate the biofilm giving neutrophils access into the infection site to improve clearance. Taken together, our results show that PNAG shelters *S. aureus* biofilms from innate host defense, and that targeting the biofilm matrix with glycoside hydrolases is a promising therapeutic avenue to treat *S. aureus* biofilm infections.

**Author Summary:** *Staphylococcus aureus* is a major cause of biofilm-associated infections, which pose a major threat to human health. A biofilm is difficult to treat since bacteria are protected from antimicrobials within an extracellular matrix. This study is the first to show that the PgaB enzyme, a glycoside hydrolase, can disrupt the *S. aureus* biofilm matrix in vivo. Disrupting the biofilm matrix with PgaB gives neutrophils access to bacteria for elimination.

## Introduction

*Staphylococcus aureus* is a leading cause of bacterial infection and foreign-body associated infections worldwide, and alternatives to antibiotics are needed to defeat these infections. The ability of *S. aureus* to form biofilm on surfaces such as catheters and medical implants, or host tissues such as bone and heart valves, contributes to its persistence in chronic infections (1). Biofilms are difficult to eradicate, as they present as a physical barrier to the host immune system and cells within the biofilm are tolerant to antimicrobials (1). The standard of practice to resolve biofilm infections is removal of the infected device, which puts the patient more at risk for developing further complications such as surgical site infections (2). Currently, there are no therapeutics in practice that are specifically designed to disrupt the *S. aureus* biofilm.

A first step of biofilm formation often involves the attachment of planktonic bacterial cells to a surface such as a foreign body, followed by growth and maturation of the biofilm (3). We previously described that pre-formed aggregates of *S. aureus* in the early stages of biofilm development were resistant to neutrophil-mediated killing and that neutrophils recruited to the skin were unable to eliminate the bacteria (4). This observation led us to hypothesize that components of the *S. aureus* biofilm matrix hinder the innate immune response leading to a persistent infection.

The extracellular matrix is a hallmark of biofilm formation. The biofilm matrix functions to protect bacteria from hostile environmental factors such as the host immune system as well as antibiotics (5). In many bacterial species, biofilm matrix polysaccharides such as cellulose, acetylated cellulose, poly-β-1,6-*N*-acetyl-D-glucosamine (PNAG), Pel, and alginate are produced by various biosynthetic machineries (6, 7). In addition to the biofilm matrix polymers, other components of the biofilm matrix include proteins, teichoic acids, and extracellular DNA (eDNA) (5).

Staphylococcal biofilms are divided into two broad categories depending on the composition of the EPS: polysaccharide-dependent and polysaccharide-independent protein-rich biofilms. Staphylococci produce one main exopolysaccharide in the EPS, initially designated as the polysaccharide intercellular adhesin (PIA), which is made up of repeating monomers of ß-1-6-linked *N*-acetyl glucosamine monomers to form PNAG. This extracellular polymer is synthesized by the enzymatic machinery encoded in the *icaADBC* operon (8, 9). Partial deacetylation of PNAG by IcaB modifies the charge of the polymer which is required for biofilm formation (10). Since its original description in *S. epidermidis* (11), the *icaADBC* operon has been identified in multiple *S. aureus* strains including *S. aureus* MW2, a clinically relevant USA400 methicillin-resistant *S. aureus* (MRSA) strain (12, 13). In addition, clinical isolates of *S. aureus* have been shown to upregulate *icaADBC* compared to commensal *S. aureus* species (14, 15). *In vivo*, PNAG has been shown to establish an invasive lung infection where deletion of *ica* in *S. aureus* resulted in fewer lung abscesses and a reduction in bacterial burden from the lungs of infected mice (16).

Due to the rise of antibiotic resistant strains, alternatives to treating *S. aureus* infections are needed. One strategy is to disrupt the biofilm by targeting the exopolysaccharide biofilm matrix with glycoside hydrolase enzymes (17). Previous studies have shown that the glycoside hydrolases α-amylase and cellulase can degrade *Pseudomonas aeruginosa* and *S. aureus* biofilms *in vitro* and *ex vivo* (18, 19). Dispersin B (DspB), which degrades PNAG (20), has been shown to reduce *S. aureus* colonization in a rabbit model of subcutaneous implant infection (21). In addition, we have previously shown that *S. aureus* biofilms can be disrupted by the glycoside hydrolase enzyme PgaB *in vitro*, where PgaB hydrolyzes partially deacetylated PNAG (22). However, no studies have determined the role of PNAG or glycoside hydrolase treatment on immune function during *S. aureus* infection *in vivo*.

In the present study, we have used a foreign body biofilm infection model using *S. aureus* agar beads to image the host response to *S. aureus* biofilm, as previously described (23). We show that the persistence is due to PIA exopolysaccharide which acts as a physical barrier that protects bacteria from the host immune system and hinders neutrophils from accessing bacteria. Infection with biofilm-deficient *S. aureus* mutants in the bead increased immune cell infiltration into the infection site at 24 hours, leading to enhanced bacterial clearance at 7 days. Imaging the biofilm revealed that neutrophils utilized elastase to infiltrate into the infection site but were sequestered at a distance from bacterial clusters separated by the PIA biofilm. After infection with biofilm-deficient strains, neutrophils were able to migrate further into the infection site and access bacterial clusters. Finally, enzymatically disrupting the biofilm matrix with PgaB, an enzyme that hydrolyzes partially deacetylated PNAG (22), improved neutrophil access to bacterial clusters and eradication in vivo suggesting that disrupting the biofilm matrix with glycoside hydrolases is a promising therapeutic option to treat biofilm infections.

## Results

Despite the evidence that *S. aureus* readily forms biofilm on foreign material such as indwelling catheters, medical devices, prostheses, and host tissue, the innate immune response to *in vivo* biofilm infection remains largely unexplored. To investigate the immune response to biofilm infection *in vivo*, we utilized a foreign body skin infection model in combination with microscopy, transgenic reporter mice and genome-engineered strains of *S. aureus* MW2, a clinically relevant MRSA strain (24).

### PIA exopolysaccharide is produced during *in vivo* biofilm infection

We have recently described a low dose foreign body infection model using *S. aureus*-contaminated agar beads (500 CFU/bead). This low dose infection formed biofilm *in vivo* and mice infected with an *S. aureus* bead took longer to clear infection compared to mice infected with planktonic *S. aureus* (23). Using this infection model, we investigated the role of polysaccharide-dependent biofilm formation by engineering an *S. aureus* MW2 strain with a complete, unmarked deletion of all 4 genes in the intercellular adhesion (*Ica*) operon (Δ*icaADBC*, see methods). At 24 hours post-infection, beads embedded with wild type *S. aureus* MW2 in beads showed evidence of biofilm formation, with clusters of *S. aureus* cells interconnected together by strands of exopolysaccharide filaments, similar to that reported previously (25) **(Figure 1A)**. By contrast, only single cells of *S. aureus* and no interconnected biofilm matrix were observed from Δ*icaADBC S. aureus* embedded beads **(Figure 1B)**. This biofilm phenotype was also validated *in vitro* using the Calgary Biofilm Device (26, 27), and the microplate biofilm assay (28) where Δ*icaADBC* formed significantly less biofilm when compared to wild type bacteria **(Figure 1C-D)**.

**Figure 1.**
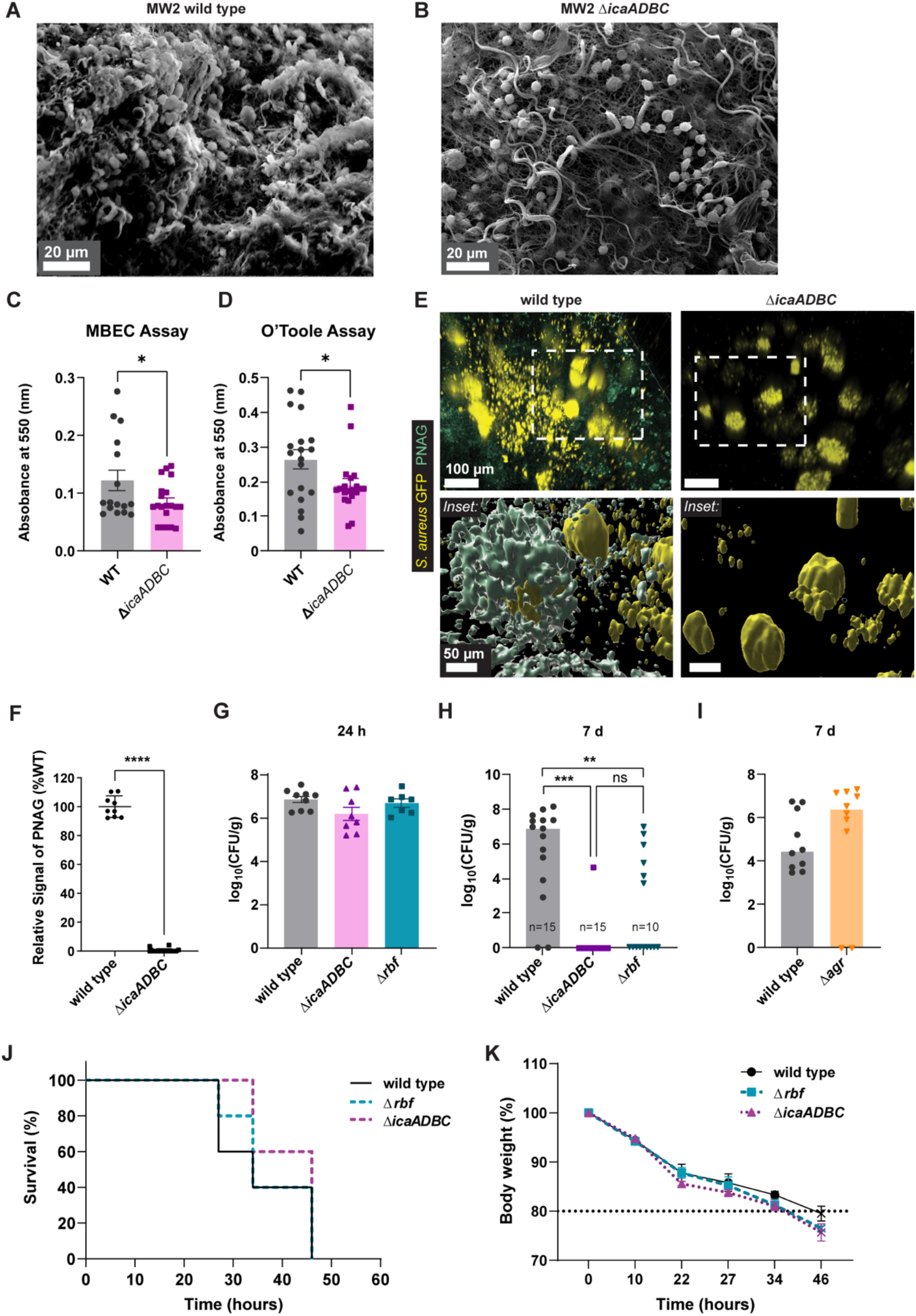
PNAG biofilm matrix contributes to *S. aureus* persistence in skin. **(A-B)** Representative scanning electron microscopy images of MW2 wild type and Δ*icaADCB S. aureus* bead at 24 hours post-infection. **(C-D)***S. aureus* MW2 wild type or Δ*icaADCB* was grown in TSB containing 0.125% glucose for 24 hours and biofilms were stained with 0.1% crystal violet using two different in vitro biofilm assays. Quantification of *in vitro* biofilm production using the MBEC assay (formerly Calgary Biofilm Device) **(C)** and O’Toole assay **(D)** assays. A 10-fold dilution was made for the O’Toole assay. *n =* 4-5 technical replicates per group from 4 independent experiments. \**p* < 0.05. Student t-test was used. **(E)** C57 mice were infected with GFP-expressing *S. aureus* wild type or Δ*icaADCB* and imaged at 24 h. Topical addition of Alexa Fluor 594-conjugated anti-F598 was added onto the skin infection prior to imaging. Image shows a 3D reconstruction (top) and surface rendering (bottom) of PNAG in wild type and Δ*icaADCB* infections. Scale bars = 100 μm (top) and 50 μm (bottom). **(F)** Semi-quantitative PNAG dot blot for *S. aureus* strains containing precisely-engineered deletions of *icaADBC* relative to the wild type strain. *n* = 3 biological and technical replicates were tested. **(G-I)** C57 mice were infected with *S. aureus* bead and infections were harvested for quantification of skin CFUs. **(G)** Bacterial CFUs at 24 hours post-infection with wild type, Δ*rbf* and Δ*icaADBC*. *n* = 7-9 from 2 independent experiments. **(H)** Bacterial CFUs at 7 days post-infection with wild type, Δ*rbf* and Δ*icaADBC*. *n* = 10-15 from 3 independent experiments. **(I)** Bacterial CFUs at 7 days post-infection with MW2 wild type and Δ*agr*. *n =* 8-10 from 2 independent experiments. **(J-K)** C57 mice were infected with 1×10^8^ CFU *S. aureus* wild type, Δ*rbf*, or Δ*icaADBC* i.v. and mouse survival **(J)** and body weight **(K)** were recorded. *n* = 5 per group from 1 independent experiment. Student t-test **(C-D, F)** and Kruskal-Wallis test (*P* = 0.0002) with Dunn’s multiple comparisons test **(H)** were used. \*\**p* < 0.01, \*\*\**p* < 0.001. \*\*\*\**p* < 0.0001.

Using multiphoton intravital microscopy, we imaged the infection at 24 hours and labeled the biofilm matrix with a fluorescently-conjugated monoclonal antibody (F598) that recognizes PNAG (29). A 3D reconstruction of *S. aureus* GFP^+^ clusters within the infection site revealed that PNAG (blue-green) was visualized surrounding wild type bacteria (yellow) in skin, which was not observed in Δ*icaADBC* infections **(Figure 1E)**. Dot blots also confirmed PNAG production *in vitro* in wild type *S. aureus* but not Δ*icaADBC S. aureus* **(Figure 1F)**.

We next tested the ability for mice to clear *S. aureus* bead infections in the presence or absence of biofilm. In addition to the Δ*icaADBC* strain, we tested a strain lacking a regulator of the *ica* operon, Rbf, which represses the negative regulator of the *ica* operon, *icaR* (13). Although there were no differences in skin CFUs between wild type, Δ*icaADBC*, or Δ*rbf S. aureus* at 24 hours post-infection **(Figure 1G)**, bacterial clearance at 7 days post-infection was dependent on *ica*-dependent biofilm formation, as both Δ*icaADBC* and Δ*rbf* bead infections were cleared from skin **(Figure 1H).** Complementation of the Δ*icaADBC* operon was not possible as trans-complementation in *S. aureus* with a useable vector in mice has not yet been developed. By contrast, *S. aureus Δagr*, which lacks *agr*-dependent quorum sensing and secreted virulence factors, did not contribute to bacterial persistence at 7 days post-infection **(Figure 1I)**.

To test the virulence of biofilm-deficient strains in a systemic challenge, we administered wild type, Δ*icaADCB* or Δ*rbf S. aureus* i.v. to C57 mice. Mortality and changes in body weight were unaffected with biofilm-deficient *S. aureus* strains after systemic infection **(Figure 1J-K)**.

### Increased immune infiltration in the absence of biofilm

In our previous work, infection with *S. aureus* embedded beads disrupted the collagen structure in the subcutaneous fascia which we visualized by second harmonic generation (SHG), and we identified the infection site by an absence of collagen signal (23). Identification of the infection site allowed us to quantify immune cell infiltration into the collagen-free zone using 3D image analysis **(Figure 2A)**.

**Figure 2.**
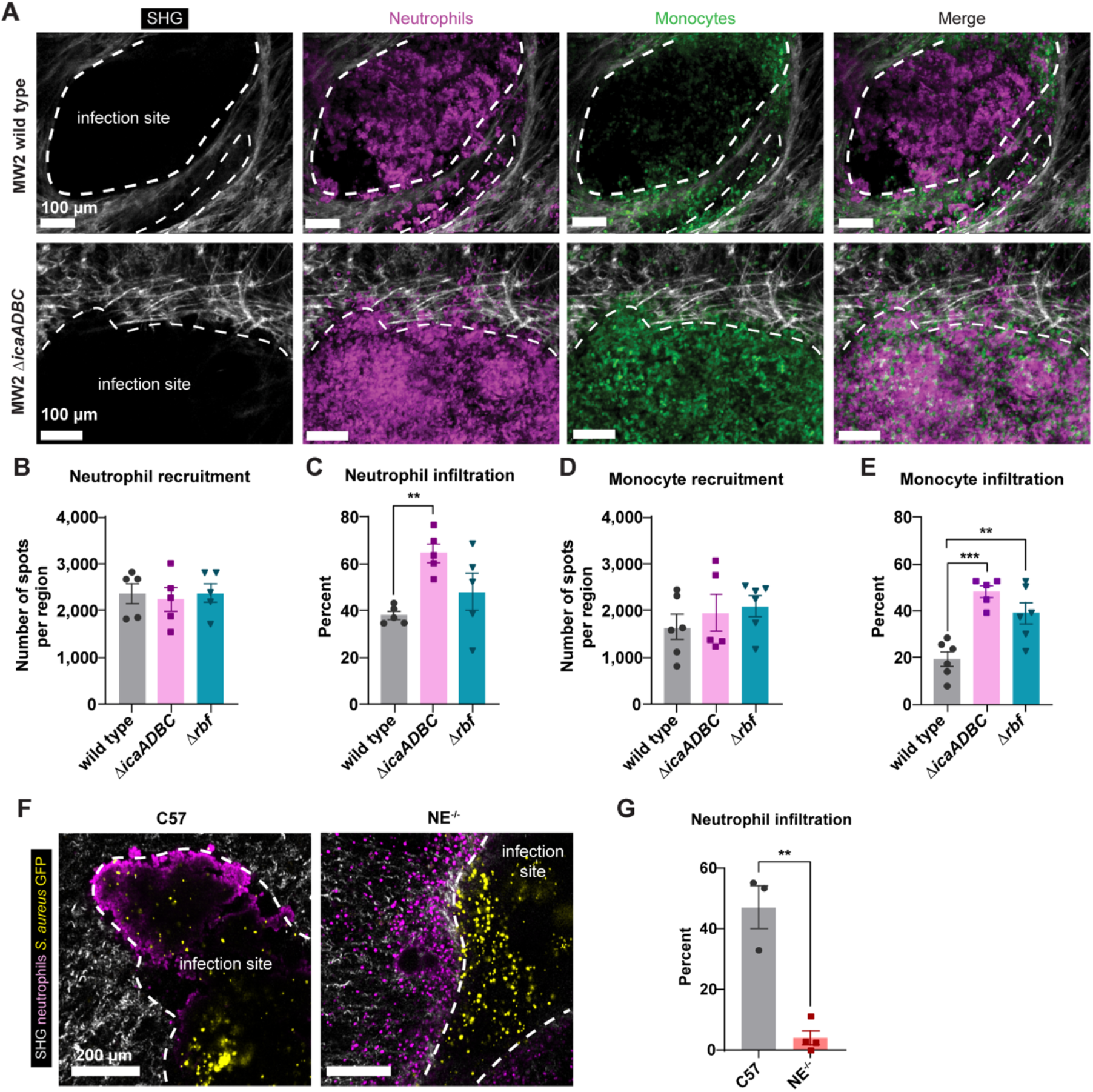
*S. aureus* biofilm blocks immune infiltration into the infection. **(A-E)** Catchup^ivm-red^ CX3CR1^gfp/wt^ mice were infected with *S. aureus* MW2 wild type, Δ*rbf*, and Δ*icaADCB* bead and imaged at 24 h. **(A)** Representative stitched image of infections in wild type and Δ*icaADCB* infected mice. Scale bars = 100 μm. **(B-E)** Quantification of total numbers of neutrophils **(B)**, percent neutrophil infiltration **(C)**, total numbers of monocytes **(D)**, and percent monocyte infiltration **(E)** at 24 hours post-infection. *n =* 4-5 per group from two independent experiments. **(F-G)** C57 or NE^-/-^ mice were infected with GFP-expressing *S. aureus* wild type and skin tissue was processed for whole mount immunofluorescence staining and imaged on the multiphoton microscope. **(F)** Representative 2D images at 24 h post-infection. **(G)** Quantification of percent neutrophil infiltration. *n* = 3-4 from 2 independent experiments. **(C, E, G)** Student t-test **(G)** and One-way ANOVA (*P* = 0.0118 for **C**, *P* = 0.0003 for **E**) with Tukey’s multiple comparison test **(C, E)** were used. \*\**p* < 0.01, \*\*\**p* < 0.001.

Although wild type and biofilm-deficient *S. aureus* bead induced similar immune cell recruitment **(Figure 2B, D)**, we observed differences in their localization pattern with respect to the collagen-free infection site. Whereas 40% of total neutrophils infiltrated in wild type infections, mice infected with Δ*icaADBC S. aureus* bead had increased neutrophil infiltration (60% of total neutrophils) **(Figure 2C)**. To see if this was just a barrier, other cell types should also be able to infiltrate into the infection site. When we looked at monocytes, they too infiltrated the infection site more effectively in the biofilm mutants than in the wild type infections suggesting that it is not specific to neutrophils **(Figure 2E)**. Neutrophil elastase (NE) regulated neutrophil infiltration into the infection site at 24 hours as NE^-/-^ mice infected with wild type *S. aureus* exhibited significantly less neutrophil recruitment into the infection site compared to C57 wildtype mice **(Figure 2F-G)**.

Flow cytometry of 24-hour skin infections confirmed that total numbers of Ly6C^int^ Ly6G^+^ neutrophils and Ly6G^-^ Ly6C^hi^ CD64^+^ monocyte-derived cells were similar among wild type, Δ*icaADCB* and Δ*rbf* strain infected mice **(Figure 3A-C)**. Additionally, there were no obvious differences in neutrophil phenotype at 24 hours post-infection in the absence of biofilm **(Figure 3D-F)**. Neutrophil behaviour was analyzed at 24 hours and after infection with Δ*icaADCB S. aureus* bead exposed neutrophils were migrating at increased velocity compared to neutrophils from wild type infected mice **(Figure 3G, Video S1)**. In the absence of biofilm, neutrophils also migrated further distances (20-40 μm) compared to neutrophils from wild type infections which migrated less than 20 μm **(Figure 3H)**. These data suggest that loss of *S. aureus* MW2 biofilm-linked traits allows neutrophils to crawl longer distances and infiltrate into the infection to encounter bacteria.

**Figure 3.**
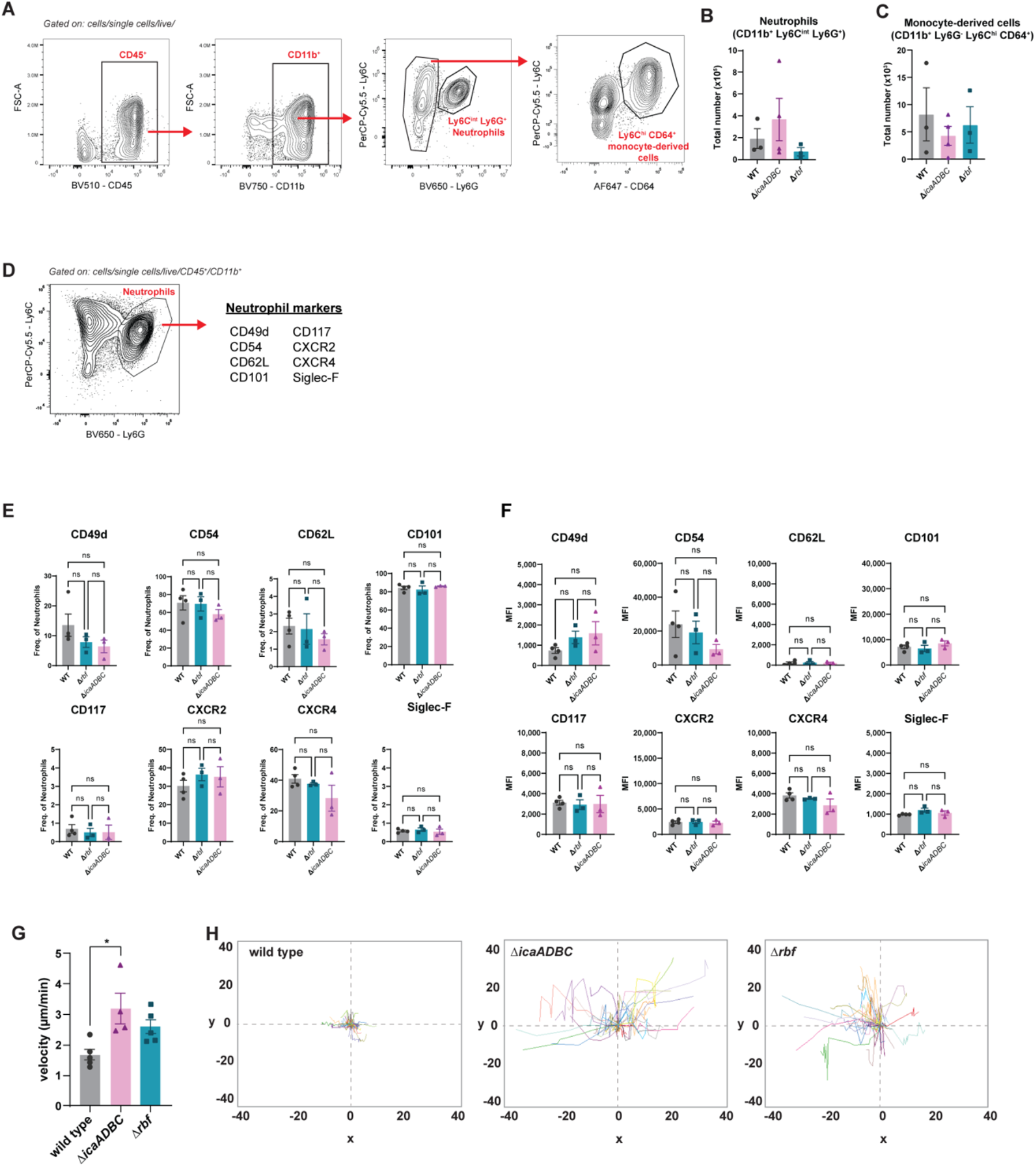
Neutrophil heterogeneity and behaviour after *S. aureus* biofilm infection. **(A-F)** C57 mice were infected *S. aureus* wild type, Δ*rbf*, and Δ*icaADCB* bead and spectral flow cytometry was performed at 24 h post-infection. **(A)** Gating strategy to identify neutrophils and monocytes. **(B-C)** Quantification of total numbers of neutrophils **(B)** and monocytes **(C)** at 24 h post-infection. **(D)** List of neutrophil markers used to characterize neutrophil heterogeneity at 24 h post-infection. Quantification of frequency **(E)** and MFI **(F)** of neutrophil subsets. *n =* 3-4 from one independent experiment. **(G-H)** Catchup^ivm-red^ mice were infected with wild type, Δ*rbf*, or Δ*icaOP S. aureus* bead and neutrophil behaviour was analyzed at 24 hours. **(G)** Quantification of neutrophil velocity over a 10-minute video. *n =* 4-5 from 2 independent experiments. **(H)** Representative spider plots showing neutrophil displacement. Each neutrophil track is identified by different colors. **(B-C, E-F)** One-way ANOVA with Tukey’s multiple comparisons test were used. **(G)** One-way ANOVA (*P* = 0.0165) with Tukey’s multiple comparison test were used. \**p* < 0.05.

### PNAG biofilm restricts neutrophil migration into the infection site to access bacteria

We hypothesized that the biofilm matrix exopolysaccharide hindered neutrophil migration into the infection site. We imaged Catchup^ivm-red^ mice infected with GFP-expressing *S. aureus* and observed a dark zone defined by the SHG-negative, GFP-negative and tdTomato-negative area **(Figure 4A)** exactly where the PNAG biofilm matrix was observed previously within the infection site in wild type infections **(Figure 1C-D)**. As we imaged deeper into the infection site below the collagen surface, the bacterial clusters became visible, and we observed a dark zone around the wild type bacterial clusters that was not apparent in biofilm-deficient infections **(Figure 4B-D)**. The dark zone area was significantly reduced in Δ*rbf* or Δ*icaADCB* infections **(Figure 4E)**.

**Figure 4.**
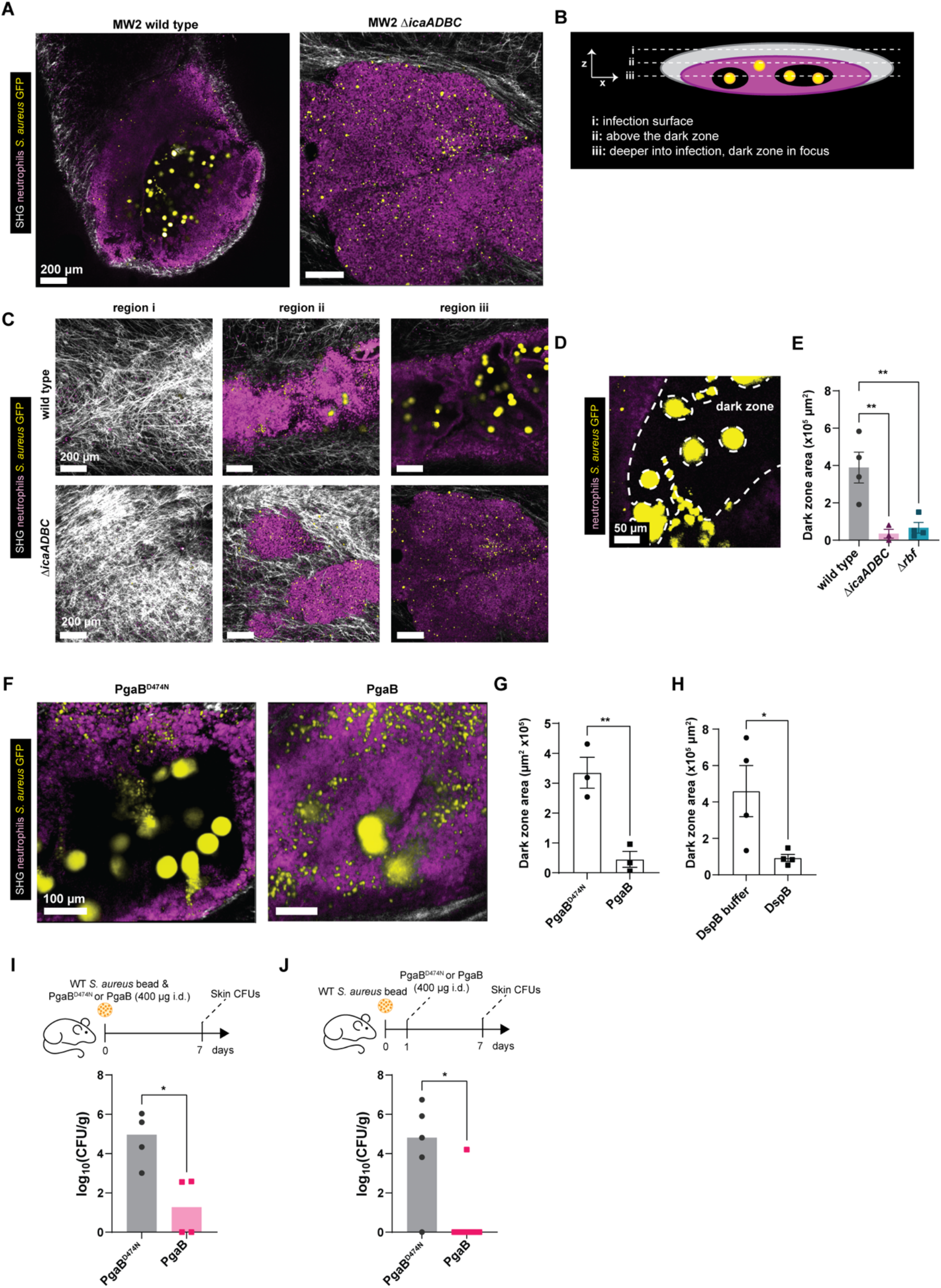
Glycoside hydrolases disrupt *S. aureus* biofilm in vivo. **(A-E)** Catchup^ivm-red^ mice were infected with GFP-expressing *S. aureus* MW2 wild type, Δ*rbf*, and Δ*icaADCB* and imaged at 24 hours. **(A)** Representative 2D images of wild type and Δ*icaADCB* infections at 24 hours. Scale bar = 200 μm. **(B)** Schematic showing 3 focal planes of a 3D z-stack of the infection. **(C)** Representative images showing 3 focal planes of a 3D z-stack in wild type and Δ*icaADCB* infections. Scale bar = 200 μm. **(D)** Representative image showing the dark zone highlighted by a dashed line. Scale bar = 50 μm. **(E)** Quantification of dark zone area in wild type, Δ*rbf*, and Δ*icaADCB* infections. *n =* 3-4 per group from 2 independent experiments. **(F-G)** Catchup^ivm-red^ mice were infected with GFP-expressing *S. aureus* wild type, treated with PgaB or PgaB^D474N^, and imaged at 24 hours. **(F)** Representative 2D images showing the dark zone after PgaB enzyme treatment. Scale bar = 100 μm. **(G)** Quantification of dark zone area in PgaB^D474N^ or PgaB treated mice. *n* = 3 per group from 2 independent experiments. **(H)** Catchup^ivm-red^ mice were infected with GFP-expressing *S. aureus* wild type, treated with DspB or DspB buffer, and imaged at 24 hours. Quantification of dark zone area in DspB or DspB buffer treated mice. *n* = 4 per group from 2 independent experiments. **(I-J)** C57 mice were infected with *S. aureus* wild type bead, treated with PgaB or PgaB^D474N^ at the time of infection **(I)** or at 24 h post-infection **(J)** and skin infections were harvested at 7 days post-infection for CFUs. *n =* 4 per group **(I)** and *n* = 5-7 per group **(J)** from 2 independent experiments. **(E, G, H-J)** One-way ANOVA with Tukey’s multiple comparison test (*P* = 0.0038) **(E)** and Student t-test **(G, H-J)** were used. \**p* < 0.05, \*\**p* < 0.01.

As we have previously shown that *S. aureus* biofilms can be disrupted by a glycoside hydrolase enzyme PgaB *in vitro* (22), we next tested whether PgaB treatment *in vivo* would affect neutrophil localization and access to bacteria. Catchup^ivm-red^ mice were infected with GFP-expressing wild type *S. aureus* embedded beads, treated with PgaB or catalytically inactive enzyme PgaB^D474N^ at the time of infection, and imaged at 24 hours post-infection. The dark zone was apparent in inactive PgaB^D474N^ treated mice but absent in active PgaB treated mice **(Figure 4F-G)**, suggesting that inhibiting biofilm production with PgaB allows neutrophils to access the bacteria. Dispersin B (DspB), another enzyme that hydrolyzes PNAG (22), also gave neutrophils access to bacteria with a reduction in dark zone area at 24 hours post-infection **(Figure 4H)**. As a result of neutrophils accessing bacteria, PgaB enzyme treatment, both at the time of infection **(Figure 4I)** or therapeutically at 24 hours post-infection **(Figure 4J)**, significantly reduced bacterial burden at 7 days post-infection.

## Discussion

These data show that the PNAG biofilm matrix produced by proteins encoded by the *icaADBC* operon is critical for *S. aureus* persistence in a foreign-body skin infection model. Imaging the early innate immune response at 24 hours post-infection revealed that immune infiltration into the infection site was hindered in the presence of biofilm which can be therapeutically targeted using the glycoside hydrolase enzyme PgaB. This work is promising for the development of new targeted therapies for foreign-body associated infections with *S. aureus*.

*S. aureus* readily forms biofilm on foreign body surfaces or host tissues. This is a major cause of chronic infections in humans (1). Despite decades of research on bacterial biofilms, most studies are limited to *in vitro* models which do not reflect what happens *in vivo* (30). Using *in vivo* intravital microscopy on living mice during a foreign body biofilm infection, we were able to image not only the biofilm *in vivo*, but also the dynamic host immune response to *in vivo* biofilm infection. Our lab had previously used a foreign-body infection model with *S. aureus*-laden agar beads but at high concentrations of bacteria (~10^6^ CFU/bead) to study neutrophil recruitment to the *S. aureus* bead (31). We have adapted this infection model to use low concentrations of *S. aureus* on the agar bead (500 CFU/bead) which serves as a foreign body and in our opinion is more clinically relevant than the more common high dose planktonic infections.

### Evidence for an *in vivo* biofilm model

Our previous data using bead infection model supported biofilm formation *in vivo* (23). We know that PIA/PNAG is present in *S. aureus* biofilm matrix which is synthesized by proteins encoded in the *icaADBC* operon (32), and many clinical isolates of *S. aureus* display high levels of *icaADBC* expression compared to commensal *S. aureus* strains (14, 15). The *S. aureus* strain used in this study, MW2, is also PIA/PNAG^+^ and forms *ica*-dependent biofilm formation *in vivo* (13). In our study, a loss-of-function mutation in *icaADBC* led to a decreased biofilm phenotype with the absence of PNAG biofilm matrix in *S. aureus*. The Δ*icaADBC* strain was cleared from skin by 7 days post-infection indicating that PIA/PNAG is critical for *S. aureus* to establish a persistent subcutaneous infection *in vivo*.

Our data with the Δ*icaADBC* strain was also supported with a *S. aureus* Δ*rbf* mutant. Rbf regulates *S. aureus* biofilm formation *in vitro* (13). A Δ*agr* mutant strain is unable to produce several virulence factors including toxins, phenol soluble modulins, and protein A (33), yet more biofilm compared to wild type strains (34). Here, deletion of *agr* in did not affect bacterial persistence at 7 days post-infection and even trended towards more difficult to eradicate infection.

In addition to PIA/PNAG, *S. aureus* can produce other biofilm matrix polymers which include teichoic acids, proteins, and eDNA. These polymers can have a variety of functions during different stages of biofilm development (5). In addition, the composition of the polymers within biofilms can vary by strain and foreign body surface (5). Future directions using this low-dose *S. aureus* bead infection model would be to identify additional components within the biofilm matrix and determine their functional role during *in vivo* biofilm infection in skin.

### Neutrophil-biofilm interactions *in vivo*

Neutrophil behaviour in close proximity to bacteria within the infection site after wild type infections and its counterparts lacking *rbf* and *icaADBC* showed striking differences despite having similar numbers of neutrophils recruited to the infected area. Neutrophils in the wild type skin infection had reduced motility whereas neutrophil in the biofilm-deficient strains were more active with higher velocity and further track displacement. This phenotype was most prominent in *S. aureus* mutant lacking *icaADBC*. The reduced motility is likely due to the presence of PIA/PNAG biofilm matrix.

Without labeling for PNAG, a dark zone that was SHG (collagen)-negative, GFP *S. aureus*-negative, and tdTomato (neutrophil)-negative was apparent which we referred to as the biofilm. The dark areas observed in *S. aureus* biofilm infections were similar to the ‘dead zone’ found after *Pseudomonas aeruginosa* biofilm infection in the eye where a black area devoid of tdTomato^+^ neutrophils and GFP bacteria was formed between host and pathogen (35). We cannot rule out the possibility of other biofilm components such as proteins or eDNA being present within the dark zone. However, after infecting with *rbf*- and *icaADBC*-deficient *S. aureus* strains, or treating mice with PgaB, the dark zone was eliminated suggesting that the PIA biofilm matrix contributed to the formation of the dark zone.

### Degradation of the biofilm *in vivo*

PgaB is a two-domain enzyme produced by the *pgaABCD* operon in *Bordetella bronchiseptica* and *Escherichia coli* species amongst others (7). Its N-terminal domain exhibits PNAG deacetylase activity while its C-terminal domain has glycoside hydrolase activity. The enzyme is required for PNAG-dependent biofilm formation (22). In staphylococcal biofilms, PgaB can disrupt the exopolysaccharide of the biofilm matrix. Importantly, pathogens who do not produce the PNAG exopolysaccharide are not affected by glycoside hydrolase activity (36).

## Conclusion

Our work adds to a growing body of work that suggests glycoside hydrolases may be used therapeutically. Although PgaB has been shown to disrupt staphylococcal biofilm *in vitro* (22), PgaB has not been used to target *in vivo* biofilms, which makes us the first to study the *in vivo* effects of this enzyme on the host immune system. In addition to Dispersin B which has the same enzymatic activity as PgaB, glycoside hydrolases from other bacteria such as *P. aeruginosa* and *Aspergillis* have also shown therapeutic potential in *in vivo* infection models (37, 38). Finally, a recent systematic survey of bacterial EPS operons revealed that 288 bacterial species contain the operon that encodes for PIA/PNAG (7), which suggests not only an evolutionarily conserved mechanism of PIA-mediated biofilm formation, but points to PgaB as a broadly targeting therapeutic for chronic infections caused by many different bacterial species. Altogether, this paper demonstrates a role for *ica*-dependent biofilm formation *in vivo* and a therapeutic angle to target chronic biofilm infections with glycoside hydrolase enzymes.

## Materials and Methods

### Mice

Animal experiments were performed with adult male and female 7-8-wk-old mice and all experimental animal protocols were approved by the University of Calgary Animal Care Committee and followed guidelines established by the Canadian Council for Animal Care (protocol number AC19-0138). All mice were housed under specific pathogen-free conditions and received sterilized rodent chow and water *ad libitum*. Mice infected longer than 24 hours were housed in a biohazard facility biosafety level 2. C57BL/6J mice were purchased from The Jackson Laboratory and bred in house. Ly6G-cre/Ai14 (Catchup^IVM-red^) mice were a kind gift from Matthias Gunzer. We crossed Catchup^IVM-red^ mice to CX3CR1^gfp/gfp^ mice to generate Catchup^IVM-red^ CX3CR1^gfp/wt^ double reporter mice, as previously described (23).

### Staphylococcus aureus

*S. aureus* strain USA400 MW2 (Baba et al., 2002) and its genetically engineered mutant strains were used for every experiment. MW2 WT and MW2 Δ*rbf* were kindly gifted to us from Dr. Chia Lee (UAMS). Bacteria were grown in Brain Heart Infusion (BHI) broth medium at 37°C while shaking at 225 rpm. When required, bacteria were transformed with pCM29 to constitutively express GFP (Pang et al., 2010). For MW2-GFP growth, chloramphenicol (10 μg/mL) was added for plasmid selection. For infection, *S. aureus* strains were sub-cultured in BHI medium without antibiotics until late exponential phase (OD_660_ 1.5), washed once with sterile PBS, and resuspended in 1 mL PBS for bead or planktonic infections.

### Standard microbiological and molecular biology methods

All strains and plasmids are listed in Table S1, and primers in Table S2. All basic microbiological and molecular procedures were executed according to standard protocols(39). DNA concentrations were measured using the A_260_/A_280_ method with a Nanodrop 2000 Spectrophotometer (ThermoFisher Scientific). Protein concentrations were determined using the Pierce 660 nm Protein Assay Reagent (ThermoFIsher Scientific), which was calibrated using bovine serum albumin standards. Genomic DNA (gDNA) isolation, plasmid preparation and DNA gel extraction were performed using nucleotide purification kits purchased from Qiagen or BioBasics. Phusion DNA polymerase and BP Clonase II were purchased from ThermoFisher Scientific. Oligonucleotide primers were purchased from Integrated DNA Technologies. Sanger sequencing was outsourced to UCDNA Services at the University of Calgary. Lysostaphin, anhydrotetracycline (ATC), chloramphenicol (CHL) and carbenicillin (CAR) were purchased from Sigma-Aldrich. Antibiotic stock solutions were corrected for activity, filter sterilized, split into 0.5 ml aliquots, and stored at −70 °C until used.

### Microbiological media and buffers

Ultrapure water was used in all buffers and media. It was prepared in-house using a Milli-Q Direct Water Purification System (Millipore-Sigma). Phosphate buffered saline (PBS) was purchased as a 20 × concentrate (Amresco). It was diluted as needed in ultrapure water and then sterilized using an autoclave. *Escherichia coli* and *S. aureus* strains were routinely propagated in lysogeny broth (LB). LB was made with 10 g tryptone, 5.0 g yeast extract and 5.0 g NaCl per 1.0 L of ultrapure water. Semi-solid LB agar in Petri dishes was prepared by adding 15.0 g/L bacteriological agar to LB prior to autoclaving. *S. aureus* was propagated, as required, in brain-heart infusion (BHI) broth, tryptic soy broth (TSB), or tryptic soy agar (TSA) purchased from BD Difco and prepared according to manufacturer’s directions in ultrapure water. As required, antibiotics and/or ATC were added as follows to media for selection during genetic manipulations: for *E. coli*, CAR at 50 μg ml^-1^, and CHL at 25 μg ml^-1^; for *S. aureus* MW2: CHL at 10 μg ml^-1^, and ATC at 0.50 μg ml^-1^. All strains were stored at −70 °C in LB containing 16.7% glycerol.

### Construction of allelic exchange vectors and *S. aureus* deletion mutants

An in-frame deletion mutation in *icaADBC* was engineered in *S. aureus* MW2 using established protocols for two-step allelic exchange(40), but with some modifications. Briefly, primer pairs oTER334/oTER345 and oTER338/oTER344 were used to amplify DNA regions upstream and downstream of *icaA* and *icaC*, respectively. These PCR products were gel purified and then fused together using established protocols(41) for splicing-by-overlap-extension (SOE) PCR with primers oTER334 and oTER344, which had been tailed with *attB1* and *attB2* sequences, respectively. The SOE-PCR product was gel purified and then recombined with pKOR1 using BP Clonase II. This reaction mixture was transformed into *E. coli* DH5α via electroporation, plated onto LB + CAR agar, and incubated overnight at 30 °C. Clones bearing the desired insert were identified via established protocols for colony PCR with M13F and M13R primers (oJJH367 and oJJH368, respectively, Table S2)(41, 42). The resulting allelic exchange vector, pKOR1 *::ΔicaABDC* (pTER139, Table S1) was verified by Sanger sequencing using M13F and M13R primers.

The pTER139 plasmid was transformed into *E. coli* DC10B, outgrown, selected, and purified using standard methods to ensure restriction modification system compatibility(43) with *S. aureus* MW2. Subsequently, 5 μg pTER139 was electroporated into *S. aureus* MW2 via established protocols(43). Merodiploids were selected on TSA + CHL after 24 h incubation at 30 °C. Subsequently, several propagations were necessary to cure the pKOR1-derived allelic exchange vector. Here, 2-3 colonies were picked from these plates and transferred to 5 mL TSB + CHL and incubated overnight at 30 °C and 250 RPM; 5 μl of this culture was then transferred to 5 mL TSB + CHL, incubated for 24 h at 42 °C and 250 RPM. This culture was serially diluted and spread onto TSA + CHL and incubated at 42 °C. Merodiploid colonies were collected with a sterile swab and transferred to 5 ml TSB and incubated overnight at 37 °C and 250 RPM, and then serially diluted and spread on TSA + ATC for counterselection. Colonies were picked from these plates and spotted on TSA and TSA + CHL. Colony PCR using phenol extraction of *S. aureus* gDNA(44) from CHL-sensitive colonies was used to identify the Δ*icaADBC* mutant using primers oTER423 and oTER382 (Table S2). The deletion mutation was confirmed via Sanger sequencing of this PCR product using the same primers, yielding *S. aureus* TER49 (Table S1).

### *S. aureus* agar bead preparation

*S. aureus*-inoculated agar beads were used to model a foreign-body infection, as previously described (23). Bacteria were resuspended in 1mL PBS, adjusted to an OD_660_ of 0.100, then diluted 10-fold. 250 μL of bacteria suspension was then added to 2.25 mL of freshly autoclaved, warm, liquid 1.5% BHI agar. The agar-bacteria mixture was slowly dropped into 40 mL ice-cold mineral oil solution containing 400 μL Tween 20 to prevent bead clumping. Gentle stirring for 15 minutes in an ice bath yielded spherical *S. aureus* agar beads. From there, beads were washed with sterile PBS by spinning in a centrifuge at 2000 rpm for 10 minutes. This wash step was repeated up to eight times to remove mineral oil coating the beads. Beads were stored at 4°C for up to two days, and fresh beads were made before every experiment. Validation of *S. aureus* bead concentrations of every bead batch were confirmed by mechanical disruption three times with a 30 gauge insulin syringe in 1 mL sterile PBS. 50 μL of the bead solution was plated onto BHI agar plates and grown overnight at 37°C for enumeration of CFUs. The average of four beads determined the bead concentration of the batch.

### Infection model – *S. aureus* agar beads

For bead infections, mice were anesthetized with isoflurane gas, back hair was shaved, and hair was chemically removed with Nair hair removal cream. Nair cream was washed off with water and skin was dried with a gauze pad. Mice were tattooed with green animal tattoo ink to permanently mark the infection site. A single bead was picked up with forceps and placed onto the bore of an 18 gauge needle connected to a syringe containing 50 μL sterile PBS, and the bead was moved back by gentle pull of the syringe which moved the bead into the needle tip. The bead was injected subcutaneously into the dorsal flank skin within the tattooed region.

### Infection model – bloodstream *S. aureus* infection

Bacterial strains were grown in 3 mL BHI overnight at 37°C while shaking. Subcultures of 150 μL bacteria in 3 mL BHI were grown for 2 hours at 37°C. 1 mL of subculture was spun down and resuspended in sterile saline and adjusted to 5×10^8^ CFU/mL. Mice were injected intravenously with 200 μL of bacteria (1×10^8^ CFU inoculum) of either *S. aureus* MW2 WT, *Δrbf* or *ΔicaADBC*. Infected mice were regularly monitored for weight and illness behaviours. Euthanasia was performed when the humane endpoint was reached (20% weight loss) or clinical signs of severe systemic illness were observed.

### Antibodies

Antibodies used for flow cytometry were purchased from eBioscience, BioLegend or BD Biosciences. Dead cells were excluded by a fixable viability dye (Ghost Red 710, Tonbo Biosciences, 1:6400). The following antibodies were used for flow cytometry:

| Reagent | Source | Identifier |
| --- | --- | --- |
| Anti-mouse CD45 BV510 (clone 30-F11) | BioLegend | Cat# 103138 |
| Anti-mouse Ly6G BV650 (clone 1A8) | BD | Cat# 740554 |
| Anti-mouse CD11b BV750 (clone M1/70) | BioLegend | Cat# 101267 |
| Anti-mouse Ly6C PerCP Cy5.5 (clone HK1.4) | BioLegend | Cat# 129012 |
| Anti-mouse CD64 Alexa Fluor 647 (clone X54-5/7.1) | BD | Cat# 558539 |
| Anti-mouse CD182 (CXCR2) BUV563 (clone V48-2310) | BD | Cat# 748681 |
| Anti-mouse CD184 (CXCR4) BUV615 (clone 2B11) | BD | Cat# 752552 |
| Anti-mouse Siglec-F BUV737 (clone 1RNM44N) | BD | Cat# 750024 |
| Anti-mouse CD117 APC (clone 2B8) | BioLegend | Cat# 105812 |
| Anti-mouse CD49d FITC (clone R1-2) | BioLegend | Cat# 103606 |
| Anti-mouse CD62L BV605 (clone C068C2) | BioLegend | Cat# 104438 |
| Anti-mouse CD54 PE (clone YN1/1.7.4) | BioLegend | Cat# 116108 |
| Anti-mouse CD101 PE-Cy7 (clone Moushi101) | Invitrogen | Cat# 12-1011-82 |

### Skin Imaging

Mice were anesthetized (ketamine (200 mg/kg) and xylazine (10 mg/kg) intraperitoneally) and a jugular catheter was inserted as previously described (McDonald et al., 2010). Anesthetics were administered through the intravenous catheter at regular intervals to maintain the mouse in a surgical plane of anesthesia. The skin surgical procedure was performed as previously described (Yipp et al., 2012). Briefly, mice were placed on a heating pad (World Precision Instruments) maintained at 37° C. A midline incision was made on the dorsal side of the mouse and the dorsal side of the flank skin was exposed and secured using silk stay-sutures onto a skin prep board (3D-printed in house). A superfusion system was set up with Hank’s Balanced Salt Solution (HBSS) heated to 37° C to perfuse the exposed skin tissue at a flow rate of 0.05. A cover glass was placed on top of the exposed skin for imaging.

### Resonant Scanning Multiphoton Intravital Microscopy

Intravital image acquisition of the skin was performed with an upright Leica SP8 resonant scanning multiphoton microscope equipped with a 20X 0.95 NA water objective. A tunable multiphoton laser was set to 940nm for excitation of GFP, tdTomato and qTracker655 for blood vessels. Second harmonic generation was visualized at an emission of 470nm. External hybrid detectors were used to detect emission at 620-680 nm (HyD-RLD1), 565-620nm (HyD-RLD2), 495-565 nm (HyD-RLD3) and <495 nm (HyD-RLD4). For 3D time lapse videos, three fields of view were selected within the infection area at 50 μm z-stack with 5 μm z-step size. Intervals were set to 30.0 seconds and videos were acquired over 20 minutes. After videos were acquired, two-three 3D regions were imaged to capture the infection area at the following dimensions: 2×5 tile scan at 200 μm z-stack with 5 μm z-step size. All videos and images used a line averaging of 16.

### Image processing, analysis and quantification

Raw imaging data was processed and quantified using Imaris Bitplane version 9.5. A Gaussian filter and background subtraction was applied to all images. A MATLAB XTension “Channel Arithmetics” was run to subtract GFP signal from (Ch2[neutrophils]-Ch3[*S. aureus* GFP]). 3D surface models of collagen were generated in Imaris using default parameters. Imaris spot function was used for automated cell counting using default parameters, and then all spots were filtered by volume to exclude spots < 5 μm^3^. For quantification of neutrophil and monocyte infiltration, a 25 μm z-stack was cropped at the focal plane of the infection site, a mask was applied to the collagen surface, then an intensity max filter was applied to neutrophil (tdTomato^+^) or monocyte (GFP^+^) spots as collagen mask^+^ or collagen mask^-^. For quantification of immune cell track length and velocity, tdTomato^+^ spots were tracked with Brownian motion over the 10-minute video and position XYZ coordinates were exported, and added into Rstudio where the data were quantified using an R script for generation of the spider plots and velocity measurements. For quantification of the dark zone in biofilm infections, a 2D region was cropped at the infection site, and the sum of collagen area, neutrophil area and GFP bacteria area was subtracted from the total region area. For overlapping areas, a mask was applied, and a new channel was duplicated such that any signal inside the mask was set to zero to ensure that the overlapping areas were not duplicated.

### CFU Experiments

Skin infections (1 cm^2^ biopsy of full thickness skin) were collected at different time points, homogenised in PBS and serial dilutions were plated on BHI agar plates and colonies were counted after 18 h at 37°C.

### Flow cytometry

Skin biopsies of the infected area were harvested (1 cm^2^ section) and collected into cold HBSS. Skin tissue was digested as previously described (23). Skin tissue was incubated at 37°C with gentle rotation for 75 minutes in 2 mL of HBSS containing 3% FBS, 5 mM EDTA and 0.8 mg/mL collagenase II (Worthington). Following enzymatic digestion tissue was passed through a 70 μm filter and washed with HBSS containing 3% FBS and 5 mM EDTA. A debris removal step was performed as per manufacturer protocol in the debris removal kit (Miltenyi Biotech). Single cells were resuspended in 800 μL of HBSS containing 3% FBS and 5 mM EDTA and 200 μL was used for antibody staining. Blood was obtained by intra-cardiac collection from anesthetized mice and 50 μL of blood lysed with ACK lysing buffer. Cells were first stained with Fc blocking antibody (1:200) and Ghost Red 710 fixable viability dye (1:6400) in HBSS for 30 minutes on ice. CXCR4 antibody staining was done at 37°C for 20 minutes. Next, cell suspensions were stained for remaining surface antigens in HBSS supplemented with 3% FBS and 5 mM EDTA for 20 minutes on ice. After washing with HBSS containing 3% FBS and 5 mM EDTA, cells were fixed with 1% paraformaldehyde in HBSS for 15 minutes on ice and then run the following day on the Cytek spectral cytometer. All flow cytometry experiments were analyzed with FlowJo v10 (Tree Star).

### In vivo enzyme treatment for biofilm disruption

A PgaB ortholog of *Bordetella bronchiseptica* (PgaB_*Bb*_) that only contained the C-terminus of PgaB was shown to have effective glycoside hydrolase activity comparable to other enzymes such as dispersin B (Little et al., 2018), and this was the enzyme used in this work. The biofilm-disrupting enzyme, PgaB, or mutant, nonfunctional enzyme PgaB D474N were injected subcutaneously into the mouse flank skin at the time of infection with the *S. aureus* bead. For PgaB and PgaB D474N, 0.4 mg/mouse was injected in a 50 μL volume.

### Whole mount 3D multiphoton microscopy

Skin samples were harvested from mice and fixed in 4% PFA for 48 hours at 4°C. Samples were then washed 3 times in 1% PBS for 1 hour at 4 °C, shaking and left in 1% BSA 1% Triton X-100 PBS overnight in 4%, shaking. Tissues were then stained with rabbit anti-S100a9 antibody (Abcam, 1:500) and chicken anti-GFP antibody (AvesLabs, 1:200) in 1% Triton X-100 and 1% BSA PBS for 3 days 4 °C, shaking. Next, the samples were washed and stained with secondary antibodies (1:1000) for 2 days 4 °C, shaking. The tissues were washed and mounted onto a glass coverslip and imaged using an Leica SP8 multiphoton microscope.

### Scanning Electron Microscopy

Skin infections were harvested and immediately fixed in 3% glutaraldehyde and paraformaldehyde for 2 hours. Samples were dehydrated in increasing concentrations of ethanol (30, 50, 70, 80, 90, and 100%), 10 minutes for each wash. Samples were transferred to hexamethyldisilazane for 1 hour and air dried overnight. Samples were sputter coated with 10nm platinum prior to imaging. SEM imaging was done on the XL30 30 kV Scanning electron microscope.

### Semi-quantitative PNAG dot blots

Dot blots for PNAG were executed by protocols modified from those previously described by our groups to measure *P. aeruginosa* PSL (Harrison et al., 2020). Here, *S. aureus* strains were grown overnight on tryptic soy agar (TSA) at 37 °C. Subsequently, single colonies were picked from the agar with a sterile loop, transferred to 2 mL tryptic soy broth (TSB) containing 1% glucose, and incubated overnight at 37 °C and 250 RPM. Cell pellets were collected from 1 mL of each culture by centrifugation (21,000 RCF for 2 min), and the supernatant discarded. The cells were suspended in 250 μL of 0.5 M EDTA (pH 8.0), and boiled at 100 °C for 60 min. These boiled suspensions were again centrifuged (21,000 RCF for 10 min), and 220 μL of supernatant was transferred to a new tube. Protein concentrations were measured for each sample using the A280 NanoDrop method and a protein standard curve (ThermoFisher Cat. 23208). Afterwards, 10 μL of proteinase K was added to each sample, and the samples were incubated at 56 °C for 30 min, followed by a 100 °C incubation for 15 min to inactivate proteinase K. These samples were diluted to 3000, 1500, and 750 ng/μL, and stored at −20 °C.

Frozen samples were thawed and then boiled at 100 °C for 5 min. A 2 μL aliquot for each sample was spotted in technical triplicate onto nitrocellulose membranes. The membrane was air dried and then rinsed in TRIS-buffered saline containing 0.5% w/v Tween-20 (TBS-T). Blots were blocked with TBS-T containing 5% w/v skim milk powder for 30 min, which were placed at room temperature on orbital shaker at 100 RPM. Afterwards, the blocking buffer was replaced with TBS-T + 5% w/v skim milk powder containing the human-α-PNAG antibody (Kelly-Quintos, Cavacini, Posner, Goldmann, & Pier, 2006) at a 1:1000 dilution. The blot was then incubated at room temperature on the orbital shaker for 1 h, and then rinsed 3 times with TBS-T (3 × 10 min, on the orbital shaker). Subsequently, the blot was labelled with horse-radish peroxidase (HRP) conjugated goat anti-human IgG antibody (Invitrogen, catalog number 31410) using a 1:3333 dilution in TBS-T. The blot was then incubated at room temperature on the orbital shaker for 1 h, and then rinsed 3 times with TBS-T (3 × 10 min, on the orbital shaker). Subsequently, the blot was labelled with horse-radish peroxidase (HRP) conjugated goat anti-human IgG antibody (Invitrogen, catalog number 31410) using a 1:3333 dilution in TBS-T. The blot was again incubated at room temperature on the orbital shaker for 1 h, and then rinsed 3 times with TBS-T as described above. Finally, the HRP-conjugated secondary antibody was visualized with Super Signal West Dura Extended Duration Substrate (ThermoFisher Scientifc^®^) and imaged using the FluorChemQ gel documentation system (Proteinsimple^®^). Images were captured and analyzed using Alphaview (v3.4.0) software (Proteinsimple^®^). Analysis was executed in biological and technical triplicate. Each sample was run in technical triplicate. One biological replicate was run per blot, 3 biological replicates were tested.

### *In vitro* biofilm assay

*S. aureus* MW2 WT or Δ*icaADCB* was grown in TSB medium containing 0.125% glucose for 24 hours and biofilms were stained with 0.1% crystal violet using two different *in vitro* biofilm assays: the Minimum Biofilm Eradication Concentration (MBEC) assay utilizing the Calgary Biofilm Device, as previously described (Ceri et al., 1999; Harrison et al., 2010) and the microplate biofilm assay (O’Toole, 2011).

### Statistical analysis and experimental design

In most experiments sample size was determined based on previous studies within the lab using these techniques. For intravital microscopy, we were limited by imaging only one mouse at a time so a minimum of 1 experimental mouse and 1 control mouse was imaged per day. Sample size was determined based on prior studies and literature using similar experimental paradigms. In instances where the approach had not previously been used, a minimum of 4 animals/group were utilized. All experiments were replicated at least once with similar findings and all replications were successful. For all experiments that required either pharmacological treatment or different infection conditions, mice were randomized. The investigators were not blinded during experiments because treatments and data collection were performed by the same researcher. For image analysis, the images were randomly assigned a key by the researcher and all images were processed using the same workflow, therefore, image analysis was blinded after data collection.

Statistical analyses were performed using Prism 9 (Graphpad Software Inc., v9.1.1, La Jolla, CA). Statistical tests are described for each figure in the figure legend. A normality test was performed for all data to determine whether a parametric or non-parametric statistical test would be used. A *P* value < 0.05 was considered statistically significant.

## Acknowledgments

The authors thank T. Nussbaumer for mice husbandry and genotyping, Dr. R. Gamutin for monitoring infected mice in the University of Calgary biohazard animal facility, Dr. P. Colarusso, Dr. A. Chojnacki, and Dr, L. Swift at the University of Calgary Live Cell Imaging Facility for microscope usage and image analysis support, Dr. J. Zindel for providing the R scripts for image analysis, Dr. W.Y. Lee for microscope assistance and maintenance and assistance with experiments, Dr. H. Kuipers for use of the Cytek Aurora which was funded through the Canadian Foundation for Innovation, Dr. P. Mukherjee and the Microscopy and Imaging Facility at the University of Calgary for scanning electron microscopy assistance, and Dr. C. Lee for providing the *S. aureus* MW2 WT and MW2 Δ*rbf* strains for our work. R.M.K. was supported by the Alberta Graduate Excellence Scholarship and the University of Calgary Doctoral Scholarship. J.C. was supported by from the Margaret Gunn Endowment for Animal Health Research. This work was supported by foundation grants from the Canadian Institute of Health Research (FDN143248 to P.K., FDN154327 to P.L.H.) and CIHR Project Grant and Tier II Canada Research Chair to J.J.H; P.L.H. was supported by the Tier I Canada Research Chair (2006-2020).

## Author contributions

R.M.K. and P.K. designed experiments. R.M.K., performed experiments. Specifically, R.M.K. performed all experiments except for T.R. who generated the Δ*icaADBC* strain and performed the dot blot experiment (Fig. 1D), J.H. performed systemic *S. aureus* infections (Fig. S2 A-B), R.S. who assisted in the generation of knockout strains, and J.C. who performed the in vitro biofilm assay (Fig. S1). R.M.K. analyzed all data. D.R. and L.H. provided the PgaB, PgaB^D474N^, and DspB enzymes, G.P. provided the F598 mAb to PNAG, D.W.M. supervised the in vitro biofilm assay, J.J.H. supervised the generation of the Δ*icaADBC* strain and provided project insight and critical review of the paper. R.M.K. wrote the paper with input from all co-authors. All authors read and approved the manuscript for submission. P.K. supervised this study.

## Competing interests

G. B. Pier is an inventor of intellectual properties [human monoclonal antibody to PNAG and PNAG vaccines] that are licensed by Brigham and Women’s Hospital to Alopexx, Inc., an entity in which GBP also holds equity. As an inventor of intellectual properties, GBP also has the right to receive a share of licensing-related income (royalties, fees) through Brigham and Women’s Hospital from Alopexx, Inc. GBP’s interests were reviewed and are managed by the Brigham and Women’s Hospital and Mass General Brigham in accordance with their conflict of interest policies.

## Supplementary videos

**Video S1. Neutrophil behaviour during biofilm infection**

Catchup^ivm-red^ mice were infected with GFP-expressing *S. aureus* MW2 WT, Δ*icaADBC*, or Δ*rbf bead* and imaged at 24h post-infection. A 3D timelapse video was taken and neutrophil behaviour was analyzed using Imaris. Video shows neutrophils (magenta), *S. aureus* GFP (yellow) and collagen (grey).

## Notes

### Competing Interest Statement

The authors have declared no competing interest.

## References

1. Costerton JW, Stewart PS, Greenberg EP. Bacterial biofilms: a common cause of persistent infections. Science. 1999;284(5418):1318–22.

2. Senthi S, Munro JT, Pitto RP. Infection in total hip replacement: meta-analysis. Int Orthop. 2011;35(2):253–60.

3. Paharik AE, Horswill AR. The staphylococcal biofilm: Adhesins, regulation, and host response. Microbiol Spectr. 2016;4(2).

4. Pettygrove BA, Kratofil RM, Alhede M, Jensen PO, Newton M, Qvortrup K, et al. Delayed neutrophil recruitment allows nascent *Staphylococcus aureus* biofilm formation and immune evasion. Biomaterials. 2021;275:120775.

5. Schilcher K, Horswill AR. Staphylococcal Biofilm Development: Structure, Regulation, and Treatment Strategies. Microbiol Mol Biol Rev. 2020;84(3).

6. Poulin MB, Kuperman LL. Regulation of biofilm exopolysaccharide production by cyclic di-guanosine monophosphate. Front Microbiol. 2021;12.

7. Bundalovic-Torma C, Whitfield GB, Marmont LS, Howell PL, Parkinson J. A systematic pipeline for classifying bacterial operons reveals the evolutionary landscape of biofilm machineries. PLoS Comput Biol. 2020;16(4):e1007721.

8. Joo H-S, Otto M. Molecular basis of in vivo biofilm formation by bacterial pathogens. Chem Biol. 2012;19(12):1503–13.

9. O’Gara JP. *ica* and beyond: biofilm mechanisms and regulation in *Staphylococcus epidermidis* and *Staphylococcus aureus*. FEMS Microbiol Immunol. 2007;270(2):179–88.

10. Pokrovskaya V, Poloczek J, Little DJ, Griffiths H, Howell PL, Nitz M. Functional Characterization of Staphylococcus epidermidis IcaB, a De-N-acetylase Important for Biofilm Formation. Biochemistry. 2013;52(32):5463–71.

11. Heilmann C, Schweitzer O, Gerke C, Vanittanakom N, Mack D, Götz F. Molecular basis of intercellular adhesion in the biofilm-forming *Staphylococcus epidermidis*. Mol Microbiol. 1996;20(5):1083–91.

12. Cramton SE, Gerke C, Schnell NF, Nichols WW, Götz F. The intercellular adhesion (*ica*) locus is present in *Staphylococcus aureus* and is required for biofilm formation. Infect Immun. 1999;67(10):5427–33.

13. Cue D, Lei MG, Luong TT, Kuechenmeister L, Dunman PM, O’Donnell S, et al. Rbf promotes biofilm formation by *Staphylococcus aureus* via repression of *icaR*, a negative regulator of *icaADBC*. J Bacteriol. 2009;191(20):6363–73.

14. Peacock SJ, Moore CE, Justice A, Kantzanou M, Story L, Mackie K, et al. Virulent combinations of adhesin and toxin genes in natural populations of *Staphylococcus aureus*. Infect Immun. 2002;70(9):4987–96.

15. Kropec A, Maira-Litran T, Jefferson KK, Grout M, Cramton SE, Gotz F, et al. Poly-N-acetylglucosamine production in *Staphylococcus aureus* is essential for virulence in murine models of systemic infection. Infect Immun. 2005;73(10):6868–76.

16. Lin MH, Shu JC, Lin LP, Chong KY, Cheng YW, Du JF, et al. Elucidating the crucial role of poly N-acetylglucosamine from Staphylococcus aureus in cellular adhesion and pathogenesis. PLoS One. 2015;10(4):e0124216.

17. Kaplan JB. Therapeutic Potential of Biofilm-Dispersing Enzymes. The International Journal of Artificial Organs. 2009;32(9):545–54.

18. Fleming D, Chahin L, Rumbaugh K. Glycoside Hydrolases Degrade Polymicrobial Bacterial Biofilms in Wounds. Antimicrob Agents Chemother. 2017;61(2).

19. Redman WK, Welch GS, Rumbaugh KP. Differential Efficacy of Glycoside Hydrolases to Disperse Biofilms. Frontiers in Cellular and Infection Microbiology. 2020;10.

20. Kaplan JB, Ragunath C, Ramasubbu N, Fine DH. Detachment of Actinobacillus actinomycetemcomitans biofilm cells by an endogenous β-hexosaminidase activity. Journal of bacteriology. 2003;185(16):4693–8.

21. Darouiche RO, Mansouri MD, Gawande PV, Madhyastha S. Antimicrobial and antibiofilm efficacy of triclosan and DispersinB^®^ combination. Journal of Antimicrobial Chemotherapy. 2009;64(1):88–93.

22. Little DJ, Pfoh R, Le Mauff F, Bamford NC, Notte C, Baker P, et al. PgaB orthologues contain a glycoside hydrolase domain that cleaves deacetylated poly-beta(1,6)-N-acetylglucosamine and can disrupt bacterial biofilms. PLoS Pathog. 2018;14(4):e1006998.

23. Kratofil RM, Shim HB, Shim R, Lee WY, Labit E, Sinha S, et al. A monocyte–leptin–angiogenesis pathway critical for repair post-infection. Nature. 2022.

24. Baba T, Takeuchi F, Kuroda M, Yuzawa H, Aoki K, Oguchi A, et al. Genome and virulence determinants of high virulence community-acquired MRSA. Lancet. 2002;359(9320):1819–27.

25. Marrie TJ, Costerton JW. Scanning and transmission electron microscopy of in situ bacterial colonization of intravenous and intraarterial catheters. Journal of Clinical Microbiology. 1984;19(5):687–93.

26. Ceri H, Olson ME, Stremick C, Read RR, Morck D, Buret A. The Calgary Biofilm Device: new technology for rapid determination of antibiotic susceptibilities of bacterial biofilms. J Clin Microbiol. 1999;37(6):1771–6.

27. Harrison JJ, Stremick CA, Turner RJ, Allan ND, Olson ME, Ceri H. Microtiter susceptibility testing of microbes growing on peg lids: a miniaturized biofilm model for high-throughput screening. Nat Protoc. 2010;5(7):1236–54.

28. O’Toole GA. Microtiter dish biofilm formation assay. J Vis Exp. 2011 (47).

29. Kelly-Quintos C, Cavacini LA, Posner MR, Goldmann D, Pier GB. Characterization of the opsonic and protective activity against *Staphylococcus aureus* of fully human monoclonal antibodies specific for the bacterial surface polysaccharide poly-N-acetylglucosamine. Infect Immun. 2006;74(5):2742–50.

30. Bjarnsholt T, Alhede M, Alhede M, Eickhardt-Sørensen SR, Moser C, Kühl M, et al. The in vivo biofilm. Trends Microbiol. 2013;21(9):466–74.

31. Harding MG, Zhang K, Conly J, Kubes P. Neutrophil crawling in capillaries; a novel immune response to *Staphylococcus aureus*. PLoS Pathog. 2014;10(10):e1004379.

32. Nguyen HTT, Nguyen TH, Otto M. The staphylococcal exopolysaccharide PIA - Biosynthesis and role in biofilm formation, colonization, and infection. Comput Struct Biotechnol J. 2020;18:3324–34.

33. Kong K-F, Vuong C, Otto M. *Staphylococcus* quorum sensing in biofilm formation and infection. Int J Med Microbiol. 2006;296(2): 133–9.

34. Vuong C, Saenz HL, Götz F, Otto M. Impact of the *agr* quorum-sensing system on adherence to polystyrene in *Staphylococcus aureus*. J Infect Dis. 2000;182(6):1688–93.

35. Thanabalasuriar A, Scott BNV, Peiseler M, Willson ME, Zeng Z, Warrener P, et al. Neutrophil extracellular traps confine *Pseudomonas aeruginosa* ocular biofilms and restrict brain iInvasion. Cell Host Microbe. 2019;25(4):526–36 e4.

36. Wang S, Breslawec AP, Alvarez E, Tyrlik M, Li C, Poulin MB. Differential Recognition of Deacetylated PNAG Oligosaccharides by a Biofilm Degrading Glycosidase. ACS Chemical Biology. 2019;14(9):1998–2005.

37. Yu S, Su T, Wu H, Liu S, Wang D, Zhao T, et al. PslG, a self-produced glycosyl hydrolase, triggers biofilm disassembly by disrupting exopolysaccharide matrix. Cell Research. 2015;25(12):1352–67.

38. Snarr BD, Baker P, Bamford NC, Sato Y, Liu H, Lehoux M, et al. Microbial glycoside hydrolases as antibiofilm agents with cross-kingdom activity. Proceedings of the National Academy of Sciences. 2017;114(27):7124–9.

39. Green MR, Sambrook J. Molecular cloning: A laboratory manual. Cold Spring Harbor, New York: Cold Spring Harbor Laboratory Press; 2012.

40. Bae T, Schneewind O. Allelic replacement in Staphylococcus aureus with inducible counter-selection. Plasmid. 2006;55(1):58–63.

41. Hmelo LR, Borlee BR, Almblad H, Love ME, Randall TE, Tseng BS, et al. Precision-engineering the *Pseudomonas aeruginosa* genome with two-step allelic exchange. Nat Protoc. 2015;10(11):1820–41.

42. Harrison JJ, Almblad H, Irie Y, Wolter DJ, Eggleston HC, Randall TE, et al. Elevated exopolysaccharide levels in *Pseudomonas aeruginosa* flagellar mutants have implications for biofilm growth and chronic infections. PLoS Genetics. 2020;16(6):e1008848.

43. Monk Ian R, Shah Ishita M, Xu M, Tan M-W, Foster Timothy J. Transforming the untransformable: Application of direct transformation to manipulate genetically *Staphylococcus aureus* and *Staphylococcus epidermidis*. mBio. 2012;3(2):e00277–11.

44. Papakyriacou H, Vaz D, Simor A, Louie M, McGavin MJ. Molecular analysis of the accessory gene regulator (agr) locus and balance of virulence factor expression in epidemic methicillin-resistant *Staphylococcus aureus*. The Journal of Infectious Diseases. 2000;181(3):990–1000.

